# *cytomapper*: an R/Bioconductor package for visualisation of highly multiplexed imaging data

**DOI:** 10.1101/2020.09.08.287516

**Authors:** Nils Eling, Nicolas Damond, Tobias Hoch, Bernd Bodenmiller

## Abstract

Highly multiplexed imaging technologies enable spatial profiling of dozens of biomarkers *in situ*. Standard data processing pipelines quantify cell-specific features and generate object segmentation masks as well as multi-channel images. Therefore, multiplexed imaging data can be visualised across two layers of information: pixel-intensities represent the spatial expression of biomarkers across an image while segmented objects visualise cellular morphology, interactions and cell phenotypes in their microenvironment.

Here we describe *cytomapper*, a computational tool that enables visualisation of pixel- and cell-level information obtained by multiplexed imaging. The package is written in the statistical programming language R, integrates with the image and single-cell analysis infrastructure of the Bioconductor project, and allows visualisation of single to hundreds of images in parallel. Using *cytomapper*, expression of multiple markers is displayed as composite images, segmentation masks are coloured based on cellular features, and selected cells can be outlined in images based on their cell type, among other functions. We illustrate the utility of *cytomapper* by analysing 100 images obtained by imaging mass cytometry from a cohort of type 1 diabetes patients and healthy individuals. In addition, *cytomapper* includes a Shiny application that allows hierarchical gating of cells based on marker expression and visualisation of selected cells in corresponding images. Together, *cytomapper* offers tools for diverse image and single-cell visualisation approaches and supports robust cell phenotyping via gating.

## INTRODUCTION

Immunohistochemistry and immunofluorescence are common approaches for visualisation of proteins in tissues^1^. Standard techniques are limited by the number of markers that can be measured simultaneously, but highly multiplexed immunohistochemistry and immunofluorescence methods have recently been developed^2–6^. Sequential staining using fluorescently-labelled antibodies allows high-resolution multiplexed imaging of tens of proteins simultaneously^7,8^. Other highly multiplexed approaches use antibodies labelled with oligonucleotides^9,10^ or metal tags^11,12^ to quantify the expression of selected proteins in cell lines and tissues.

One of the latter approaches is imaging mass cytometry (IMC), a highly multiplexed imaging technique that simultaneously measures the spatial expression of up to 40 proteins (also referred to as markers) at 1μm resolution. During IMC, tissue slices are stained with metal-conjugated antibodies prior to laser-ablation and ion detection^11^. After data acquisition, raw output files are processed to create multi-channel images where pixel-intensities represent the measured ion counts per marker. These images are further processed to create segmentation masks containing object identifiers corresponding to each identified cell. Segmentation masks can be used to extract cell-specific measurements such as mean ion counts per marker and morphological features^13^.

Custom scripts^4,11,14,15^ and image analysis software such as *ImageJ*^16^, *CellProfiler*^17^, *ilastik*^18^ and *QuPath*^19^ are commonly used to process and analyse multiplexed imaging data. More recently, specialised tools based on graphical user interfaces (GUIs) have been developed to facilitate the analysis of high-dimensional spatial expression data^20–23^. The GUI toolbox *histoCAT* was specifically designed for IMC data to visualise multi-channel images and segmentation masks, and to link cellular phenotypes to their location within tissues^24^. Furthermore, commercial software developed by Visiopharm, Fluidigm (*MCD viewer*), IONpath (*MIBItracker*), Perkin Elmer (*inForm*), Indica Labs (*Halo*), TissueGnostics (*StrataQuest*) and Leica (*ImageScope*) are available to perform multiplexed image analysis^1^. While GUI-based tools facilitate data analysis for scientists with little experience of scripting languages, they are limited to a chosen set of algorithms. With the advent of single-cell RNA sequencing and mass cytometry technologies, a multitude of user-friendly analysis tools^25–27^ have been developed to perform flexible single-cell data analysis using the statistical programming language R^28^.

Here, we combine the image and single-cell data analysis capabilities of Bioconductor^29^ to allow visualisation of pixel- and cell-level information obtained by highly multiplexed imaging technologies such as IMC. The R/Bioconductor package *cytomapper* builds upon the *SingleCellExperiment* data container^27^ and uses *EBImage*^30^ functionality to visualise segmentation masks and multi-channel images. Therefore, *cytomapper* allows high flexibility in terms of image manipulation (e.g. transformations) and integrates with common single-cell data analysis strategies (e.g. cell phenotyping). The *cytomapper* package further includes a Shiny application to enable hierarchical gating of cells based on their marker expression and visualisation of selected cells in corresponding images. We demonstrate the utility of *cytomapper* by using it for biological exploration of type 1 diabetes progression and quality control of image segmentation results.

## RESULTS

The *cytomapper* package enables visualisation of pixel- and cell-level information obtained by highly multiplexed imaging technologies. The main functions of the package require a data object that stores cell-specific expression and metadata information, and data objects that contain either segmentation masks or multi-channel images.

### Technical details and implementation

Single-cell expression values and cell-specific metadata such as cell type information are stored in a *SingleCellExperiment* class object^27^ (**Fig. 1A**). The *cytomapper* package provides the *CytoImageList* container that stores single- or multi-channel images of various sizes (**Methods** and **Fig. 1A, B**). These objects can contain segmentation masks represented as single-channel images; or multi-channel images where each channel contains pixel-intensities of an individual marker. By providing information regarding a cell’s object identifier and a unique image name, the *plotCells* function colours segmentation masks by marker expression or cell-specific metadata (**Fig. 1A**). Multi-channel images are visualised as composites of up to 6 channels using the *plotPixels* function (**Fig. 1B**). Furthermore, by providing a *SingleCellExperiment* object and a *CytoImageList* object containing segmentation masks, individual cells can be outlined on composite images.

**Figure 1:**
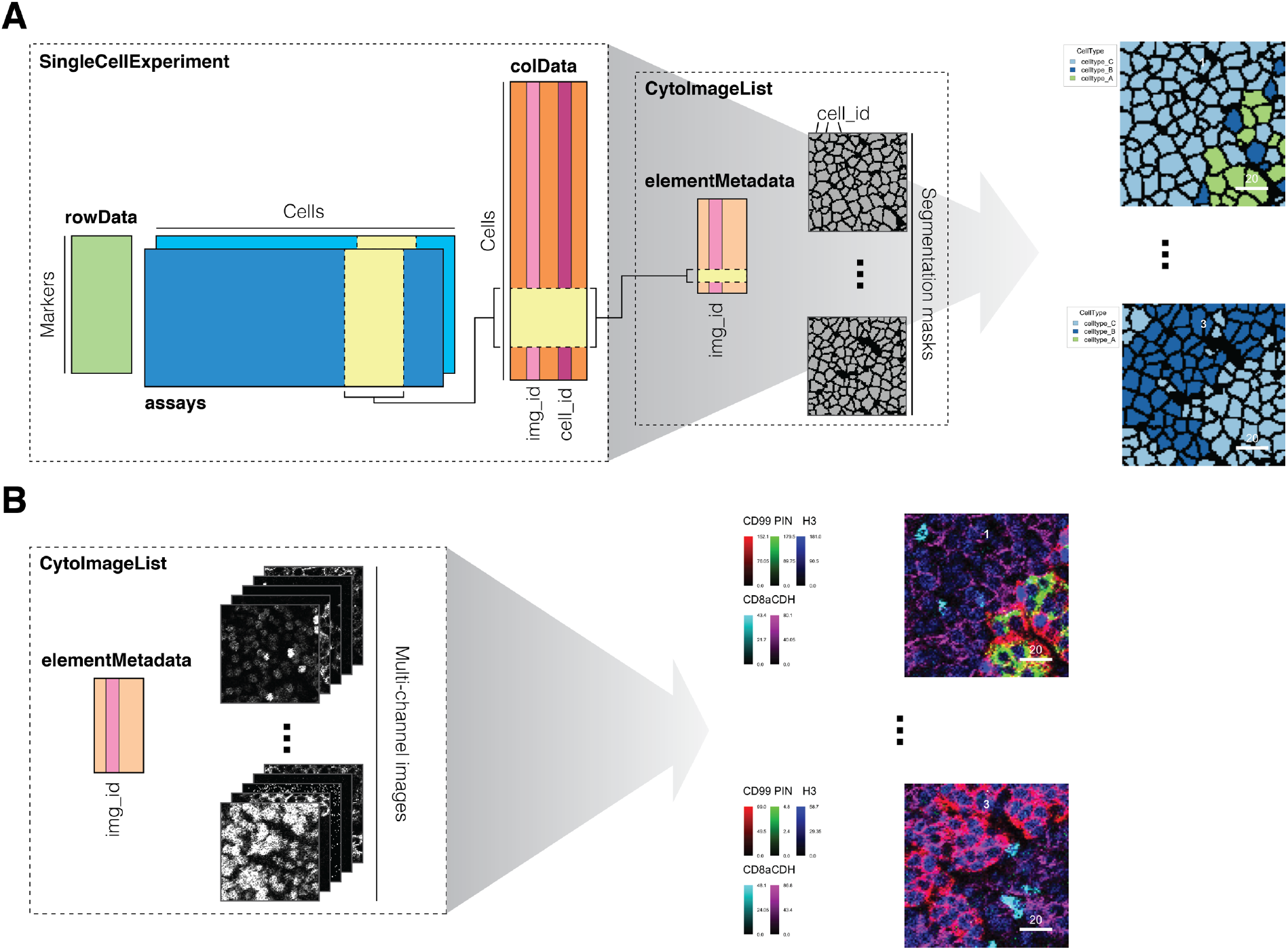
Technical implementation of the *cytomapper* package. **(A)** Single-cell expression and metadata is stored in a *SingleCellExperiment* object. Important entries are *img_id*, which stores unique image names and *cell_id*, which stores the cell object identifiers. The *img_id* entry is linked to metadata stored in the *CytoImageList* object containing segmentation masks. The *plotCells* function combines these two objects to visualise marker expression or cell-specific metadata on segmentation masks. **(B)** The *plotPixels* function requires a *CytoImageList* object containing multi-channel images to visualise the combined expression of up to six markers as composite images. In addition, a *SingleCellExperiment* object and *CytoImageList* object containing segmentation masks can be provided to outline cells and colour outlines by cell-specific metadata. The data presented here are a toy dataset provided by the *cytomapper* package. Scale bars: 20μm

### Visualising pancreatic cell types during type 1 diabetes progression

To demonstrate the functionality of the *cytomapper* package we used it to visualise type 1 diabetes (T1D) samples acquired by 35-plex IMC^13^ (**Methods**). T1D is characterised by β cell loss caused by autoreactive immune cell infiltration^31^ and we previously imaged pancreatic samples from patients with recent-onset and long-duration, as well as healthy controls^13^. We ranked images based on the density of cytotoxic and helper T cells and selected the image with highest density per condition. Using the *cytomapper* package, we visualised all islet cell types, and cytotoxic and helper T cells in selected images (**Fig. 2A**). To visually confirm cell phenotypes, we further displayed cell type specific markers (proinsulin (PIN): β cells; CD4: helper T cells; CD8a: cytotoxic T cells) on segmentation masks (**Fig. 2B**) and as composite images (**Fig. 2C**). By visualising selected images, we observe, as expected, that (i) β cells and proinsulin expression are lost during T1D progression and (ii) T cells invade the microenvironment during early onset of T1D^13^.

**Figure 2:**
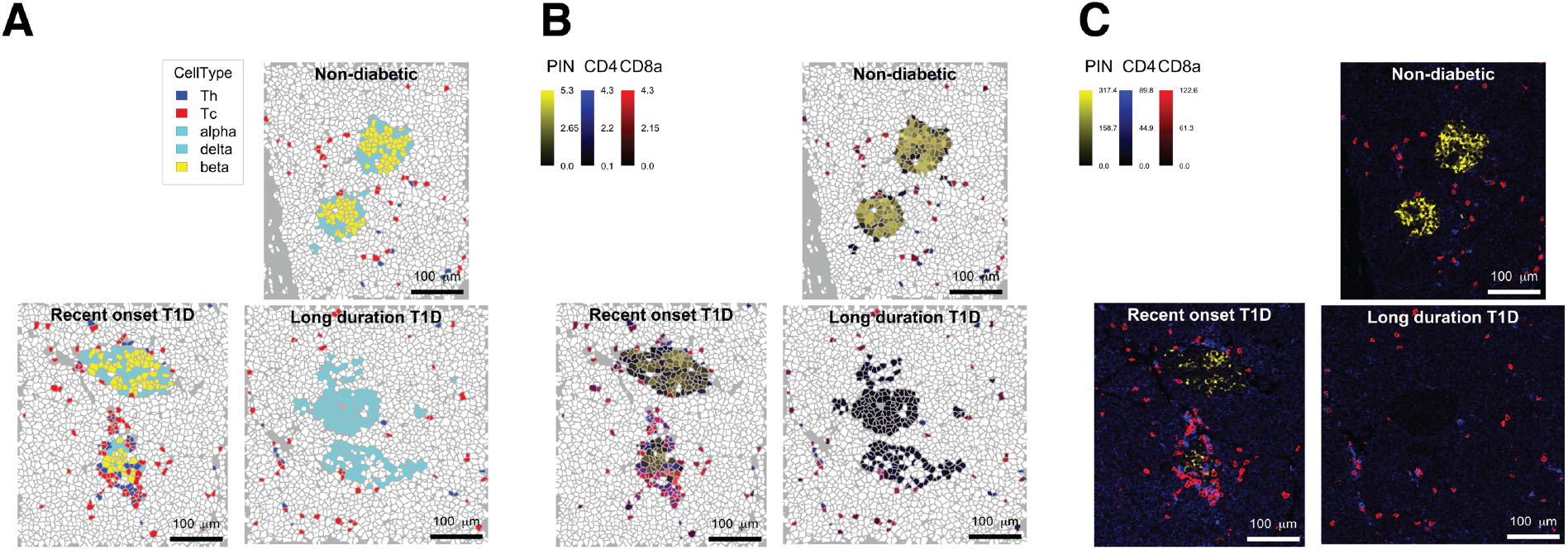
Islet and immune cell dynamics during T1D progression. For each condition (healthy, recent onset and long-duration T1D), images with the highest density of cytotoxic and helper T cells were selected. **(A)** The *SingleCellExperiment* object was subsetted to only contain islet cells, and cytotoxic and helper T cells allowing the selective visualization of those cell types. The *plotCells* function colours selected cells by their cell type and leaves all other cells white. **(B)** The selected cells are coloured based on the arsinh-transformed expression of proinsulin (PIN) in yellow, CD4 in blue and CD8a in red marking β cells, helper T cells and cytotoxic T cells respectively. **(C)** These markers are visualised as composite images by merging pixel-level information of marker expression. Raw pixel-intensities were multiplied by 10, 8 and 10 for PIN, CD4 and CD8a, respectively to increase the contrast of the images. Scale bar: 100μm

### β cell and proinsulin loss during T1D progression

The analysis performed above highlights the use of *cytomapper* to visualise a small subset of images. To avoid a biased representation of biological phenomena, *cytomapper* also allows the visualisation of tens to hundreds of images in parallel. Next, we will highlight the loss of β cells and reduction of proinsulin expression across 100 selected images from the full set of 845 images acquired to profile T1D progression^13^.

We first ranked the images based on the percentage of β cells out of all islet cells and then used the *plotCells* function to visualise islet cell types across all segmentation masks. By labelling the masks based on T1D stage (healthy, recent onset and long-duration T1D), we observe a loss of β cells along disease progression (**Fig. 3**).

**Figure 3:**
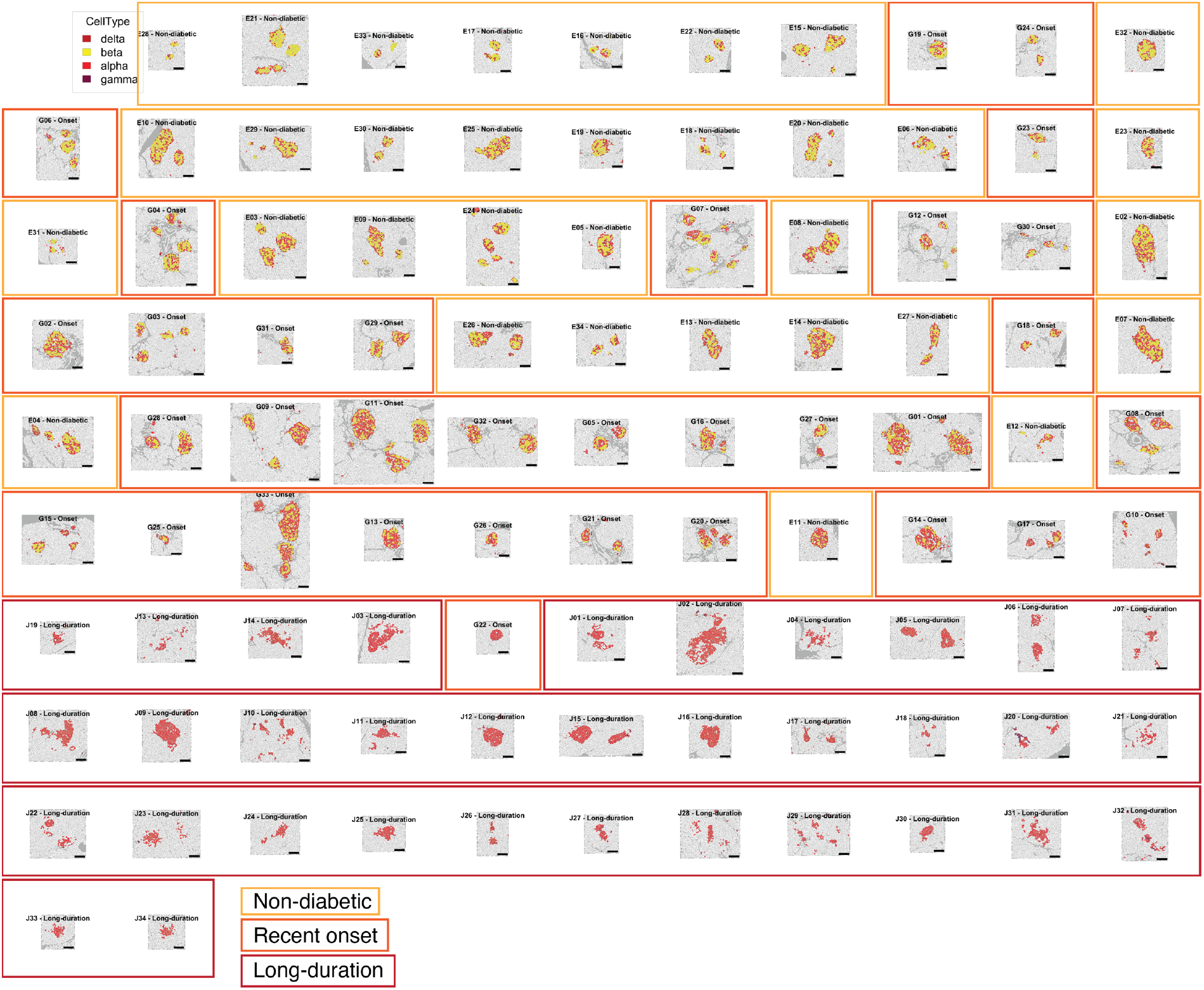
Distribution of islet cell types along T1D progression. A set of 100 segmentation masks was ordered based on the frequency of β cells out of all islet cells. We selected islet cell types for visualisation; all other cells are displayed in white. The *plotCells* function was used to colour cell areas based on their cell type. Segmentation masks are automatically arranged in a grid-like pattern supporting differences in image dimensions. Image borders are manually coloured by T1D stage (healthy, recent onset and long-duration). A progressive loss of β cells (coloured in yellow) can be observed. Scale bar: 100μm

To validate these results, we ranked multi-channel images based on the mean signal of proinsulin expression (mean pixel-intensity of pixels with a detectable signal, i.e. a signal > 0 counts). We normalised images in a two-step process. First, we performed a min-max scaling to normalise pixel intensities to 0 and 1 across all images. Next, we clipped normalised pixel intensities to 0 and 0.05 removing pixels with high outlying intensities. This approach allows the qualitative comparison of pixel intensities across images. Observing image-to-image differences in total pixel intensities can indicate batch effects in staining efficiency^21^ or biological features such as the expected loss of proinsulin signal over T1D progression (**Fig. 4**).

**Figure 4:**
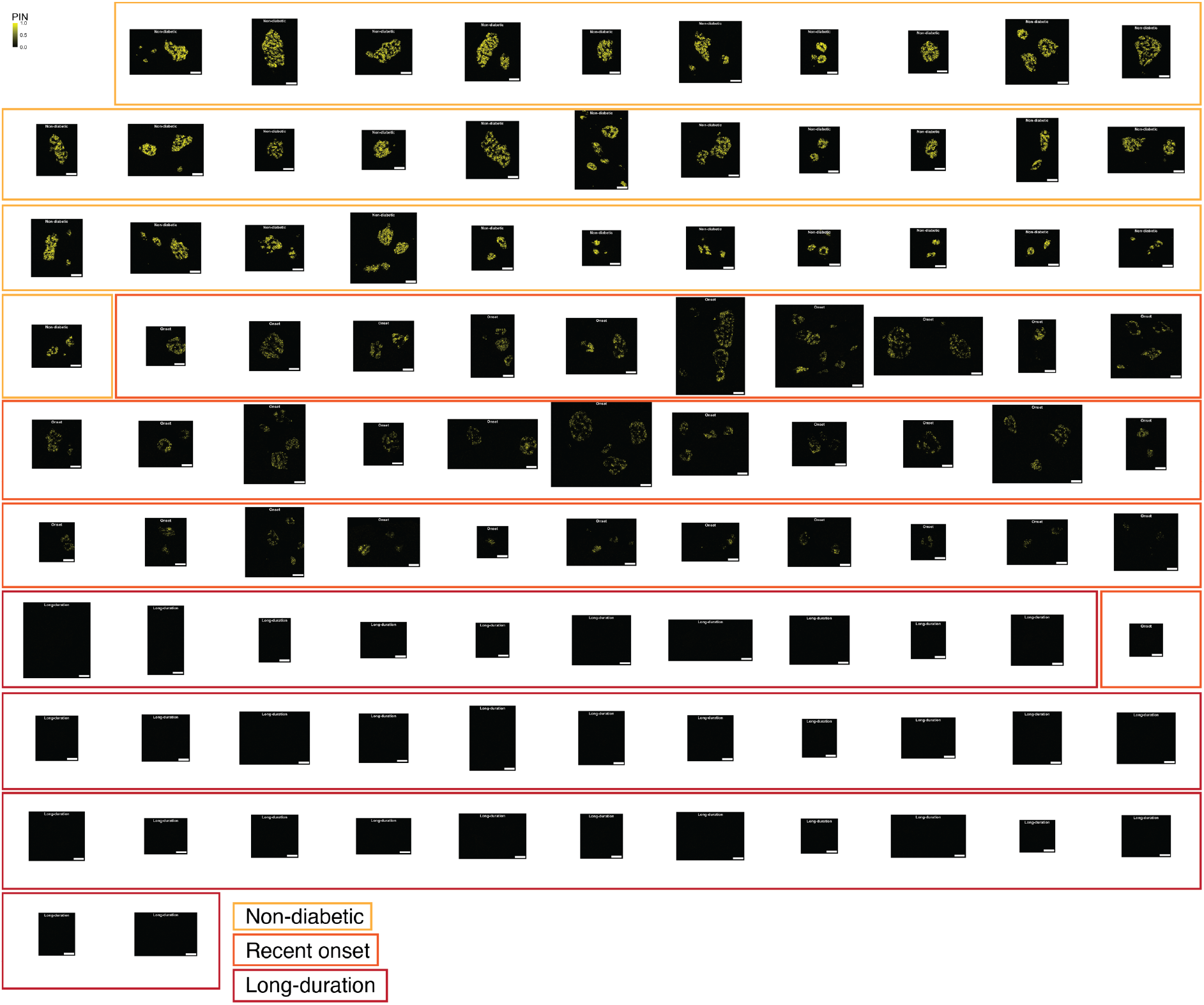
Proinsulin expression across T1D progression. A set of 100 multi-channel images was ordered by calculating the mean of all pixels with detectable proinsulin signal. Images were normalised by first scaling pixel intensities between the lowest and highest pixel intensity across all images. In the second normalisation step, normalised pixel intensities were clipped at 0 and 0.05. Image borders are manually coloured by T1D stage (healthy, recent onset and long-duration). Scale bar: 100μm

### Visual quality control of segmentation and cell-phenotyping results

Segmentation and labelling of cell phenotypes are essential steps of most multiplexed imaging pipelines^32^. The *cytomapper* package provides function settings to outline cells on composite images based on their segmentation results. Furthermore, outlines can be coloured based on cell-specific metadata, such as cell type information. Figure 5 presents results of the *plotPixels* function, where we selected specific cell types by subsetting the *SingleCellExperiment* object and provide segmentation masks. In that way, the segmentation results (shape of the outline) and cell-labelling results (colour of outline) can be compared to the spatial expression of individual or multiple markers. We detect good but not perfect overlap between outlines and marker expression (**Fig. 5**), highlighting the challenge of image segmentation and cell type detection. This visual quality control step is recommended prior to downstream analyses such as clustering or the testing of associations with clinical data.

**Figure 5:**
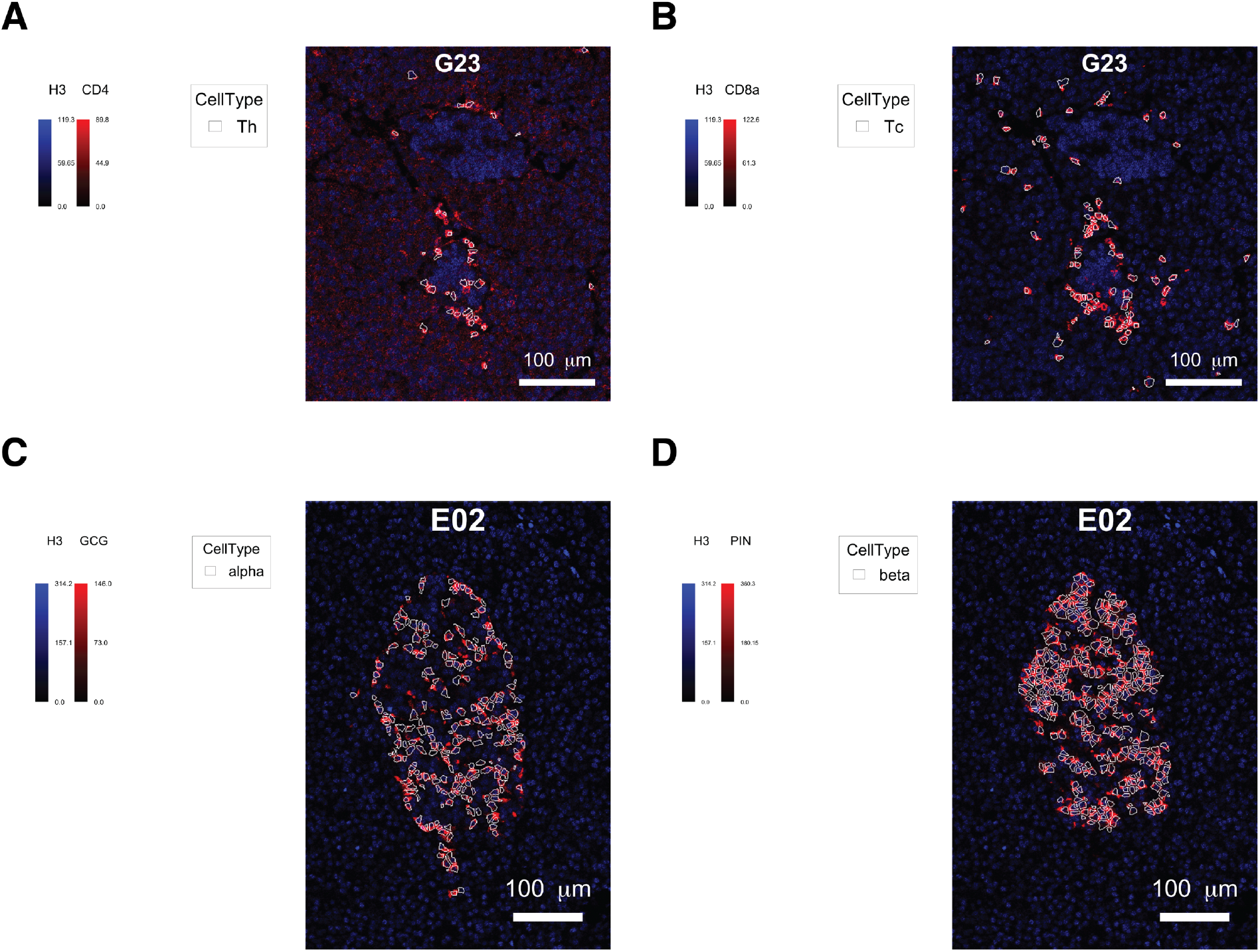
Visual quality control of segmentation and cell-labelling results. **(A) – (B)** The image with highest T cell density (helper and cytotoxic T cells) was selected. The marker H3 indicates nuclear stain, **(A)** CD4 expression marks helper T cells (Th) and **(B)** CD8a expression marks cytotoxic T cells (Tc). Individual channels were multiplied by a constant to increase the contrast (H3: 1.5, CD4: 6, CD8a: 6). The *SingleCellExperiment* was subsetted to only contain helper T cells **(A)** or cytotoxic T cells **(B)**. Cells are outlined in white based on their cell type. **(C) – (D)** All images of healthy donors were ranked based on their β cell or α cell density. The image with the lowest rank sum was selected for visualisation. The marker glucagon (GCG) indicates α cells **(C)** while proinsulin (PIN) is expressed in β cells **(D)**. Individual channels were multiplied by a constant to increase the contrast (H3: 6, GCG: 6, PIN: 6). The *SingleCellExperiment* was subsetted to only contain α cells **(C)** or β cells **(D)**. Cells are outlined in white based on their cell type.

### A Shiny application for gating and visualization of cells

Cell phenotyping is commonly performed by clustering and cluster annotation. However, a number of classification strategies have recently been developed to label cells based on a given reference^33^. In the case of highly multiplexed imaging data, images are usually annotated (e.g. nucleus, cytoplasm, background) on a pixel-level using tools such as *ilastik*^13,18^. Alternatively, cells can be labelled based on their averaged pixel intensity.

To facilitate cell labelling, we developed the *cytomapperShiny* function, which opens a *shiny* GUI that allows hierarchical gating on the expression levels of up to 24 markers. Gating is performed on the raw or transformed expression counts stored in any *assay* slot of the *SingleCellExperiment* (**Fig. 6A**). Selected cells are either visualised as coloured objects on segmentation masks (**Fig. 6B**) or as outlines on composite images (**Fig. 6C**). Furthermore, selected cells can be downloaded in form of a *SingleCellExperiment* object and for use in downstream processes such as training and cell type classification.

**Figure 6:**
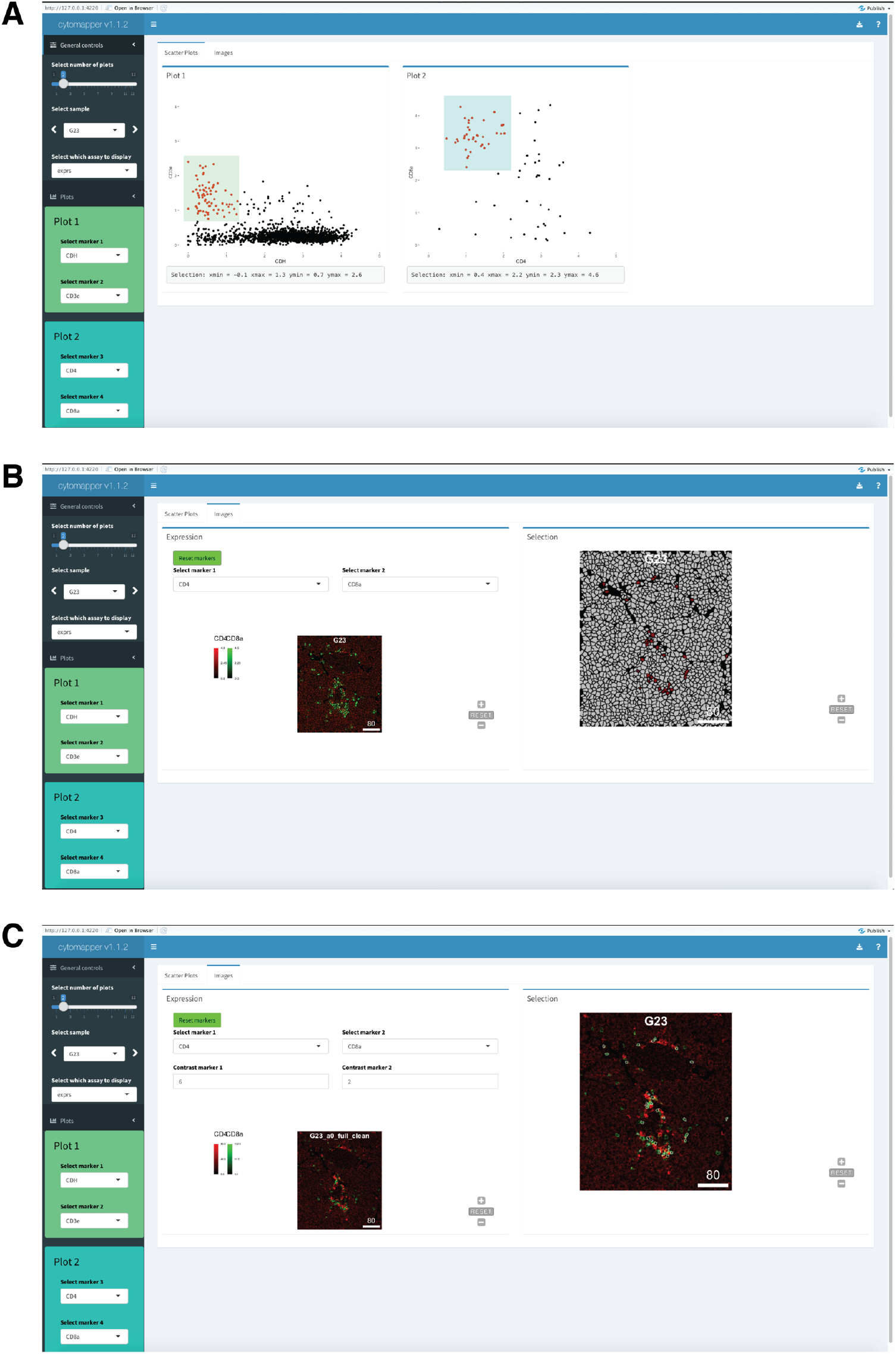
The *cytomapperShiny* GUI. **(A)** The shiny GUI provided by *cytomapper* allows gating of cells based on their raw or transformed expression counts. **(B)** If only segmentation masks are supplied, selected cells are coloured as filled objects on their corresponding segmentation mask. **(C)** If the user provides multi-channel images, selected cells are outlined on their corresponding composite image. Here, we first gated cells based on low E-cadherin (CDH) and high CD3e expression to select T cells. These cells were further sub-gated for high CD8a and low CD4 expression to select cytotoxic T cells.

## DISCUSSION

The *cytomapper* package offers a set of functions to visualise cell- and pixel-level information obtained using highly multiplexed imaging technologies. By combining the *SingleCellExperiment* container with the newly developed *CytoImageList* class object, segmentation masks are coloured based on cell-specific features. Furthermore, pixel-intensities from multiple channels are merged to be displayed as composite images.

We demonstrated the use of *cytomapper* with IMC data. However, data obtained using other multiplexed imaging technologies such as MIBI^12^, 4i^8^, CODEX^9^ and t-CyCIF^7^ and spatial transcriptomics methods including MERFISH^34^ and seqFISH^35^ could be visualised using the *cytomapper* package. The only requirements are single-cell read-outs, multi-channel tiff stacks and/or segmentation masks.

The *cytomapper* package is built upon *EBImage*^30^, which offers image analysis functionality in R. Furthermore, by using the *SingleCellExperiment* object as a data container, *cytomapper* integrates with an extensive set of single-cell data analysis tools as well as other R packages designed for spatial data analysis^23,36^.

We have demonstrated some use cases for the *plotCells* and *plotPixels* function to visualise the cellular heterogeneity within images. By subsetting the *SingleCellExperiment* object, *cytomapper* supports selected visualisation of specific cell types within tissues. This approach can be of interest when confirming segmentation and cell labelling results. Furthermore, the specific interaction of two or more cell types can be highlighted by first selecting the cells of interest.

One key advantage of *cytomapper* is the visualisation of tens to hundreds of images in parallel. The grid-like visualisation is not restricted to uniform image dimensions and signal intensities can be qualitatively compared across images. This approach is crucial to identify technical or biological staining differences between images, batches or conditions^21^.

The *cytomapper* package further offers a shiny GUI by calling the *cytomapperShiny* function. The application is based on a hierarchical gating strategy, which is widely used in flow cytometry. We extended this approach with an additional layer of information by visualising imaging data and displaying gated cells. This allows to easily monitor the result of the applied gating strategy and increases the quality of labelled data. The *shiny* application provided by *cytomapper* is unique as it combines hierarchical gating with image visualisation and integrates seamlessly with the Bioconductor infrastructure for single-cell analyses. The ease of generating and improved quality of training data will meet the growing demand for supervised classification methods^33^.

## METHODS

### Type 1 Diabetes dataset

We highlight the functionality of *cytomapper* by re-analysing a dataset acquired by imaging mass cytometry. We previously profiled 1,581 pancreatic islets from 12 donors, 4 of which were non-diabetic, 4 had recently been diagnosed with T1D (< 0.5 years) and 4 had long-standing T1D (> 8 years)^13^. For this, we developed an antibody panel consisting of 35 markers, 2 nuclear stains and PD-1, which was excluded from analysis in the original paper, to profile the interactions between immune and pancreatic islet cells during disease progression. A total of 845 images were acquired where each image captures an individual or few pancreatic islets.

Cells and pancreatic islets were segmented using *CellProfiler*^37^ and *ilastik*^18^ by selecting informative channels (https://github.com/BodenmillerGroup/ImcSegmentationPipeline). Multi-channel tiff files were directly created using the *imctools* python package (https://github.com/BodenmillerGroup/imctools)^13^. *Ilastik* was further used to classify cells into four categories: islet, immune, exocrine and “other”. Islet cells were further classified into α, β, δ and γ cells; immune cells were classified into B cells, cytotoxic and helper T cells, monocytes/macrophages, and neutrophils; exocrine cells were sub-divided into acinar and ductal cells and “other” cells were classified into endothelial, stroma and unknown^13^.

As an example dataset, we selected one representative donor per disease stage (non-diabetic, recent onset, long-duration) and randomly selected 100 images (33 non-diabetic, 33 recent-onset, 34 long-duration) from these three donors, containing a total of 252,059 cells.

A *SingleCellExperiment* object was created to store cell-specific expression values and metadata. Mean pixel intensities per cell and marker were loaded into the *counts assay* slot and arsinh-transformed counts were stored in the *exprs assay* slot. Cell-specific (e.g. cell identifier, cell type information, location, area, islet- and neighbour relationships), patient (e.g. case identifier, disease stage, age, gender) and image (e.g. image identifier, slide identifier, dimensions) metadata were stored in the *colData* entry of the *SingleCellExperiment* object. Marker-specific metadata such as metal isotope, marker name, stock concentration and channel identifier were loaded into the *rowData* slot. Attention needs to be paid to the correct ordering of the channels and cells in the *SingleCellExperiment* object.

We used the *loadImages* function provided by *cytomapper* to read in the multi-channel tiff stacks and segmentation masks into *CytoImageList* objects. The correct image identifier was added to the metadata entry of both *CytoImageList* objects to link them to information stored in the *SingleCellExperiment* object. Furthermore, the *CytoImageList* object storing segmentation masks was scaled by a factor of 65535 to account for 16-bit scaling.

### The CytoImageList container

The *CytoImageList* container provides an S4 class object to store multiple single- or multi-channel images. We created helper functions to get and set the channel names for multi-channel images and provide a number of functions for consistent subsetting. In places where segmentation masks are needed for visualisation, *cytomapper* will test if the supplied *CytoImageList* object only stores single-channel images that contain integer values or 0. The integer values indicate the numeric identifier of each cell while 0 labels the background of the image. Furthermore, the *CytoImageList* container stores metadata for each image (e.g. disease type) and supports easy subsetting. When using the *cytomapper* functionality, unique image identifiers must be provided in the *CytoImageList* metadata (see below).

The *CytoImageList* class supports storing images with different x- and y-dimensions in individual slots. However, each image needs to have the same z-dimension (same number of channels).

For convenience, we have created the *loadImages* function that reads a single image or multiple images (in .tiff, .png or .jpeg format) into R while creating a *CytoImageList* object. The user can further provide a *pattern* argument to selectively read in a subset of images.

### The SingleCellExperiment container

The *SingleCellExperiment* container is a popular S4 class used in Bioconductor workflows and tools to store information of individual cells^27^. The main slots of the object store raw and transformed expression data (*assays*), cell-specific metadata (*colData*), gene/marker-specific metadata (*rowData*) and low-dimensional embeddings (*reducedDims*). Here, we store the mean pixel intensities per cell in the *counts* assay slot and the arsinh-transformed (using a co-factor of 1) mean pixel intensities in the *exprs* assay slot. The *cytomapper* package combines *CytoImageList* and *SingleCellExperiment* objects to visualise information contained in the *SingleCellExperiment* object on images contained in the *CytoImageList* object.

### Linking images and cell-specific data

As explained above, the *CytoImageList* object contains multiple single- or multi-channel images. The unique image IDs need to be stored within the metadata of the object. In the case of segmentation masks, integer pixel values represent the cells’ object identifiers. This information is used to link cells and images to data stored in the *SingleCellExperiment* object. Both, the unique image ID and cell ID, are stored in the cell-specific *colData* slot of the *SingleCellExperiment* object and matching to the images stored in the *CytoImageList* object is done internally. In that way, the *SingleCellExperiment* object can be subsetted to only contain a specific selection of cells (see **Fig. 2**).

### The plotCells function

The *plotCells* function visualises cell-specific marker expression or metadata on segmentation masks. The main input to the *plotCells* function is a *CytoImageList* object containing the segmentation masks, a *SingleCellExperiment* object containing the cell-specific information, and the image ID and cell ID slot entries. Furthermore, the user can specify which markers or metadata to visualise, chose the colour scale per marker or metadata entry and which expression data slot to use (raw or transformed counts).

The combined expression of up to six markers can be visualised on the segmentation masks using additive colour mixing. However, we do not recommend displaying multiple markers with overlapping expression patterns on the segmentation masks. The user can specify two or more colours for each marker that are interpolated to generate a marker-specific colour scale. When displaying marker expression on segmentation masks, colours are scaled between the minimum and maximum expression count across all cells contained in the *SingleCellExperiment* object. This is also true when subsetting images prior to plotting.

Furthermore, individual metadata entries such as cell phenotype can be visualised on the segmentation mask using automatic or manual colouring. Cells can either be filled or outlined by metadata entries.

### The plotPixels function

The *plotPixels* function visualises marker expression by displaying a pseudo-colour representation of pixel-intensities. When visualising single channels, the *viridis* colour scale is used to display low intensities as blue and high intensities as yellow. Between two and six channels can be additively merged to display a pseudo-colour composite image. The default colours for displaying high intensities are red, green, blue, cyan, magenta and yellow. However, colours can be changed by providing at least two colours per channel (minimum and maximum) to generate a continuous colour scale. Colours are scaled between the minimum and maximum pixel-intensity across all displayed images. Therefore, when subsetting images before plotting, the range of pixel-intensities can change.

The user can control the brightness (b), contrast (c) and gamma value (g) of the displayed image by setting the *bcg* parameter. The *bcg* parameters used for the current figures are listed in the figure legends. Only the contrast parameter was set in the presented analyses.

Furthermore, the user can provide an additional *SingleCellExperiment* and *CytoImageList* object containing segmentation masks (see above). By doing so, cells can be outlined based on metadata features stored in the *SingleCellExperiment* object (see **Fig. 5**).

### Image normalisation

The *EBImage* R/Bioconductor package^30^ provides a *normalize* function that scales pixel-intensities between 0 and 1 either channel-wise or across all channels. The user can further provide a clipping range to set pixel-intensities to either 0 or 1 if outside of the range. We have adapted this scaling normalization to multiple multi-channel images. By default, the *cytomapper normalize* function scales pixel intensities channel-wise across all images contained in the *CytoImageList* object. This default setting is chosen to display staining differences between images^21^. The user can also choose to perform the scaling normalisation per image by setting *separateImages = TRUE*. We further provide a *scaleImages* function that multiplies the pixel intensities per image with a constant value. This is useful when read-in pixel intensities are not correctly scaled.

### Additional plotting parameters

The user can modify different features of the displayed images, save the images or get them returned in R for further analysis. These options are shared between the *plotPixels* and *plotCells* function and are documented under the key *plotting-param* in R. The colour of cells on the segmentation masks that are not contained in the *SingleCellExperiment* object can be set using the *missing_colour* parameter. The background colour can be changed by setting *background_colour*. The length, label, size, colour and position of the scale bar can be changed using the *scale_bar* parameter. Image titles can be controlled by setting the *image_title* parameter. All features of the colour legend can be controlled by setting the *legend* parameter. To save the displayed images, the user can specify the *save_plot* parameter. This takes a list containing the *filename* and a *scale* scalar x. The later scales the resolution of the image x fold. By setting *return_plot = TRUE* the displayed images including image titles and scale bars are returned as a single plot or list of plots. When setting *return_images = TRUE* a list of individual images is returned. However, scale bars and image titles are lost when returning composite images. By default, multiple images are plotted on a grid with varying margins between individual images depending on the maximum image width and height. To further increase the margin between individual images, the *margin* parameter can be set. The user can further set *display = “single”* to plot individual images in their own graphics device instead of on a grid. By default, each channel is scaled between its minimum and maximum before creating the composite image. This behaviour can be supressed and relative differences between channels can be observed by setting *scale = FALSE*. By default, pseudo-colours are interpolated between neighbouring pixels to smooth the image. Interpolation is supressed by setting *interpolate = FALSE*.

To suppress the display of the legend, image title and scale bar, their corresponding parameters can be set to *NULL*.

### The Shiny application

We developed an interactive application using the R packages *shiny* and *shinydashboard* to gate cells based on their expression values and to visualise selected cells on images. The *cytomapperShiny* function takes a *SingleCellExperiment* object storing cell-specific features and *CytoImageList* objects storing segmentation masks or multi-channel images as input. Upon execution, the function opens a graphical user interface with two tabs. In the first tab, hierarchical gating can be performed on expression values stored in the *SingleCellExperiment* object. If the user further provides segmentation masks and (optional) multi-channel images, *cytomapperShiny* visualises expression values and gated cells on images in the second tab. By using the R packages *svglite* and *svgPanZoom*, the *cytomapper* image output is converted to a scalable vector graphic which enables pan and zoom functionality. Finally, the user can download gated cells in form of a *SingleCellExperiment* object. For reproducibility purposes, *cytomapperShiny* stores the gates, the gating date and the session information in the *metadata* slot of the downloaded *SingleCellExperiment* object.

## CODE AND DATA AVAILABILITY

All analysis was performed using Bioconductor 3.12, R version 4.0.2 and *cytomapper* version 1.1.2 (available from Github with the tag v1.1.2).

All analysis code and instructions for data analysis are available at: https://github.com/BodenmillerGroup/cytomapper_publication

The exact scripts for the current manuscript have been deposited on Zenodo with the DOI 10.5281/zenodo.3994630 version v1.0

A static website visualising the results can be found at: https://bodenmillergroup.github.io/cytomapper_publication/

A docker container running the exact software used for the analysis can be obtained from: https://hub.docker.com/r/nilseling/bioconductor_cytomapper/tags tag 0.0.1

The release version of the *cytomapper* package (version 1.0.0) can be installed via Bioconductor: https://www.bioconductor.org/packages/release/bioc/html/cytomapper.html

The development version of the *cytomapper* package (version 1.1.2 or higher) containing the *cytomapperShiny* application is available via: https://www.bioconductor.org/packages/devel/bioc/html/cytomapper.html or https://github.com/BodenmillerGroup/cytomapper

The full dataset and the smaller example dataset used in the present publication are available at: https://data.mendeley.com/datasets/cydmwsfztj/2

## AUTHOR CONTRIBUTIONS

N.D. developed the first version of the software. N.E. and N.D. implemented the *cytomapper* Bioconductor package. N.E and T.H implemented the Shiny application. N.E., N.D., T.H. and B.B. wrote the manuscript.

## ACKNOWLEDGEMENTS

We want to thank Daniel Schulz and Jana Fischer for critical feedback on the package and Natalie de Souza for feedback on the manuscript.

This work was supported by the European Molecular Biology Organisation [ALTF 1194-2019]; the Juvenile Diabetes Research Foundation [3-PDF-2020-937-A-N]; and a National Institute of Health grant [DK108132].

## Notes

### Competing Interest Statement

The authors have declared no competing interest.

## REFERENCES

1. Finotello, F., Rieder, D., And, H. H. & Trajanoski, Z. Next-generation computational tools for interrogating cancer immunity. Nat. Rev. Genet. 20, 724–746 (2019).

2. Schubert, W. et al. Analyzing proteome topology and function by automated multidimensional fluorescence microscopy. Nat. Biotechnol. 24, 1270–1278 (2006).

3. Stack, E. C., Wang, C., Roman, K. A. & Hoyt, C. C. Multiplexed immunohistochemistry, imaging, and quantitation: A review, with an assessment of Tyramide signal amplification, multispectral imaging and multiplex analysis. Methods 70, 46–58 (2014).

4. Gerdes, M. J. et al. Highly multiplexed single-cell analysis of formalinfixed, paraffin-embedded cancer tissue. Proc. Natl. Acad. Sci. U. S. A. 110, 11982–11987 (2013).

5. Huang, W., Hennrick, K. & Drew, S. A colorful future of quantitative pathology: Validation of Vectra technology using chromogenic multiplexed immunohistochemistry and prostate tissue microarrays. Hum. Pathol. 44, 29–38 (2013).

6. Tsujikawa, T. et al. Quantitative Multiplex Immunohistochemistry Reveals Myeloid-Inflamed Tumor-Immune Complexity Associated with Poor Prognosis. Cell Rep. 19, 203–217 (2017).

7. Lin, J. R. et al. Highly multiplexed immunofluorescence imaging of human tissues and tumors using t-CyCIF and conventional optical microscopes. Elife 7, e31657 (2018).

8. Gut, G., Herrmann, M. D. & Pelkmans, L. Multiplexed protein maps link subcellular organization to cellular states. Science (80-.). 361, eaar7042 (2018).

9. Goltsev, Y. et al. Deep Profiling of Mouse Splenic Architecture with CODEX Multiplexed Imaging. Cell 174, 968–981 (2018).

10. Saka, S. K. et al. Immuno-SABER enables highly multiplexed and amplified protein imaging in tissues. Nat. Biotechnol. 37, 1080–1090 (2019).

11. Giesen, C. et al. Highly multiplexed imaging of tumor tissues with subcellular resolution by mass cytometry. Nat. Methods 11, 417–422 (2014).

12. Angelo, M. et al. Multiplexed ion beam imaging of human breast tumors. Nat. Med. 20, 436–442 (2014).

13. Damond, N. et al. A Map of Human Type 1 Diabetes Progression by Imaging Mass Cytometry. Cell Metab. 29, 755-768.e5 (2019).

14. Keren, L. et al. A Structured Tumor-Immune Microenvironment in Triple Negative Breast Cancer Revealed by Multiplexed Ion Beam Imaging. Cell 174, 1373-1387.e19 (2018).

15. Jackson, H. W. et al. The single-cell pathology landscape of breast cancer. Nature 578, 615–620 (2020).

16. Schneider, C. A., Rasband, W. S. & Eliceiri, K. W. NIH Image to ImageJ: 25 years of image analysis. Nat. Methods 9, 671–675 (2012).

17. Carpenter, A. E. et al. CellProfiler: image analysis software for identifying and quantifying cell phenotypes. Genome Biol. 7, R100 (2006).

18. Sommer, C., Straehle, C., Kothe, U. & Hamprecht, F. A. Ilastik: Interactive learning and segmentation toolkit. in IEEE International Symposium on Biomedical Imaging: From Nano to Macro 230–233 (2011). doi: 10.1109/ISBI.2011.5872394

19. Bankhead, P. et al. QuPath: Open source software for digital pathology image analysis. Sci. Rep. 7, 16878 (2017).

20. Stoltzfus, C. R. et al. CytoMAP: A Spatial Analysis Toolbox Reveals Features of Myeloid Cell Organization in Lymphoid Tissues. Cell Rep. 31, 107523 (2020).

21. Somarakis, A., Van Unen, V., Koning, F., Lelieveldt, B. P. F. & Hollt, T. ImaCytE: Visual Exploration of Cellular Microenvironments for Imaging Mass Cytometry Data. IEEE Trans. Vis. Comput. Graph. 1–1 (2019). doi: 10.1109/TVCG.2019.2931299

22. Czech, E., Aksoy, B. A., Aksoy, P. & Hammerbacher, J. Cytokit: a single-cell analysis toolkit for high dimensional fluorescent microscopy imaging. BMC Bioinformatics 20, (2019).

23. Dries, R. et al. Giotto, a pipeline for integrative analysis and visualization of single-cell spatial transcriptomic data. bioRxiv (2019). doi: https://doi.org/10.1101/701680

24. Schapiro, D. et al. HistoCAT: Analysis of cell phenotypes and interactions in multiplex image cytometry data. Nat. Methods 14, 873–876 (2017).

25. Ellis, B. et al. FlowCore: Basic structures for flow cytometry data. R package version 2.0.1 (2020).

26. Chevrier, S. et al. Compensation of Signal Spillover in Suspension and Imaging Mass Cytometry. Cell Syst. 6, 612-620.e5 (2018).

27. Amezquita, R. A. et al. Orchestrating single-cell analysis with Bioconductor. Nat. Methods 17, 137–145 (2020).

28. R Core Team. R: A Language and Environment for Statistical Computing. (2020).

29. Gentleman, R. C. et al. Bioconductor: open software development for computational biology and bioinformatics. Genome Biol. 5, (2004).

30. Pau, G., Fuchs, F., Sklyar, O., Boutros, M. & Huber, W. EBImage-an R package for image processing with applications to cellular phenotypes. Bioinformatics 26, 979–981 (2010).

31. Atkinson, M. A., Eisenbarth, G. S. & Michels, A. W. Type 1 diabetes. Lancet 383, 69–82 (2014).

32. Moen, E. et al. Deep learning for cellular image analysis. Nat. Methods 16, 1233–1246 (2019).

33. Abdelaal, T. et al. A comparison of automatic cell identification methods for single-cell RNA sequencing data. Genome Biol. 20, 1–19 (2019).

34. Chen, K. H., Boettiger, A. N., Moffitt, J. R., Wang, S. & Zhuang, X. Spatially resolved, highly multiplexed RNA profiling in single cells. Science (80-.). 348, aaa6090 (2015).

35. Lubeck, E., Coskun, A. F., Zhiyentayev, T., Ahmad, M. & Cai, L. Single-cell in situ RNA profiling by sequential hybridization. Nat. Methods 11, 360–361 (2014).

36. Yang, T. et al. SPIAT : An R package for the Spatial Image Analysis of Cells in Tissues. biorx 1–16 (2020).

37. Kamentsky, L. et al. Improved structure, function and compatibility for cellprofiler: Modular high-throughput image analysis software. Bioinformatics 27, 1179–1180 (2011).

